# ISCA1 and CRY4: An improbable proposition

**DOI:** 10.1101/094458

**Authors:** Tobias Hochstoeger, Simon Nimpf, David Anthony Keays

## Abstract

This manuscript was submitted to Nature Materials on December 15th 2015.

In their recent manuscript Xie and colleagues argue that the pigeon ISCA1/CRY4 complex acts as a magnetic protein biocompass having unambiguously proved that these two proteins interact forming a complex with unique biophysical features (1). We do not think the evidence presented in the manuscript supports this conclusion. Firstly, the authors provide no data (such as a reciprocal co-immunoprecipitation from retinal tissue) that demonstrates that ISCA1 and CRY4 interact *in vivo*, instead the conclusions are based primarily on overexpression in *E.coli* and *in vitro* reconstitution experiments. Secondly, their experiments have not employed the correct full length version of CRY4. The CRY4 protein expressed by Xie and colleagues was only 497 amino acids in length. We extracted mRNA from the pigeon retina (Invitrogen, 610.11), generated cDNA (Clontech, 121311), and performed rapid amplification of cDNA ends using a proof reading DNA polymerase (Thermo Scientific, #F-549S). This allowed us to clone the full length version of pigeon CRY4, which is 525 amino acids in length (Supplementary Figure 1). This peptide sequence shares a high level of homology with chicken CRY4 which is 529 amino acids in length and the zebrafinch CRY4 which is 527 amino acids long (2) (Supplementary Figure 2). We confirmed that our sequence is correct by cloning CRY4 from a second pigeon strain originating from Frankfurt. Given that the C-terminal region of CRY4 is known to be functionally important, undergoing structural change in response to light, it is unclear how to interpret the results presented by Xie and colleagues (3). Thirdly, in support of their claim that ISCA1 and CRY4 interact they performed immunohistochemistry on the pigeon retina. Employing sera raised against "full” length CRY4 and ISCA1 they report co-localization in the ganglion cell layer, the inner nuclear layer, and the outer nuclear layer of the retina. Putting aside the fact there is no data supporting the assertion that their CRY4 antibody is specific, this experiment does not demonstrate a direct interaction between the two proteins. Nor is this result remotely surprising as current evidence suggests ISCA1 is ubiquitously expressed in eukaryotes, as it plays an important role in mitochondrial biogenesis and function (4, 5). We investigated the expression of *Isca1* and *Cry4* in the pigeon by extracting total RNA from a range of organs (n=3 birds) (Qiagen, 74104), generated cDNA (Qiagen, 205314), and performed real time quantitative PCR (qPCR) employing exon spanning primers (Biorad, 1708880). We normalised our results to three control genes *(Hprt, Gapdh, Tfrc).* We made sure that the primers we used were specific by undertaking BLAST searches against the pigeon genome and transcriptome, and by analysing DNA melt curves (Table S1). We find that *Isca1* is expressed at moderate levels in all major organs with lowest expression in the retina. *Cry4* was likewise expressed in all tissues analysed, with highest levels in the beak skin, and lowest levels in the heart (Figure 1). We confirmed these results by performing qPCR with a second set of primers for *Isca1* and *Cry4* (data not shown).

**Figure 1.**
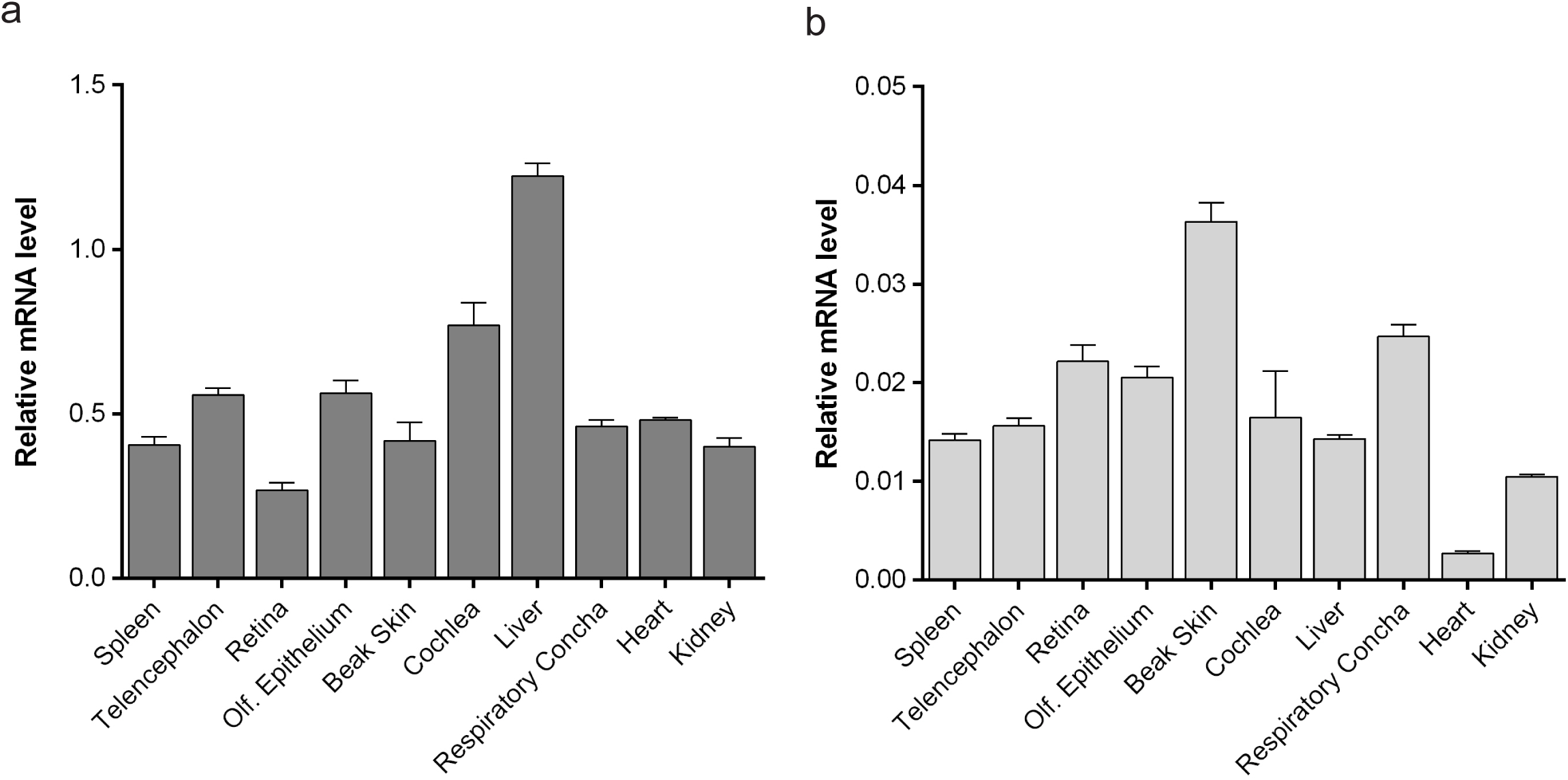
*Isca1* and *Cry4* are expressed broadly in the pigeon. (a) qPCR results showing the expression of *Isca1* in the spleen, telencephalon, retina, olfactory epithelium, beak skin, cochlea, liver, respiratory concha, heart and kidney. Expression levels were normalised to the geometric mean of three control genes (*Hprt*, *Gapdh*, *Tfrc*). Isca1 is expressed at moderate levels in all tissues with the lowest levels of expression in the retina. (b) *Cry4* is expressed in all tissues analysed, with highest levels of expression in the beak skin, and lowest levels in the heart. Error bars show standard error of the mean.

Previous studies have highlighted that molecules specialized for sensory transduction show restricted expression patterns. For instance, olfactory receptors are enriched in the olfactory epithelium (6), rhodopsin is found almost exclusively in photoreceptive cells (7), and the mechanosensitive TMC1 channel is concentrated in hair cells (8). This specialisation is further reflected in the processing of sensory information in the central nervous system where distinct anatomical regions have been associated with specific senses (e.g. visual cortex). Neuroanatomical studies in the pigeon have highlighted the importance of the vestibular and trigeminal nuclei in processing magnetic information (9, 10). In light of this fact and the observation that ISCA1 and CRY4 are expressed broadly in all adult organs of pigeons is it really conceivable that they are the primary magnetic sensors? We think this is an improbable proposition.

## Supplementary Material

**Supplementary Figure 1.**
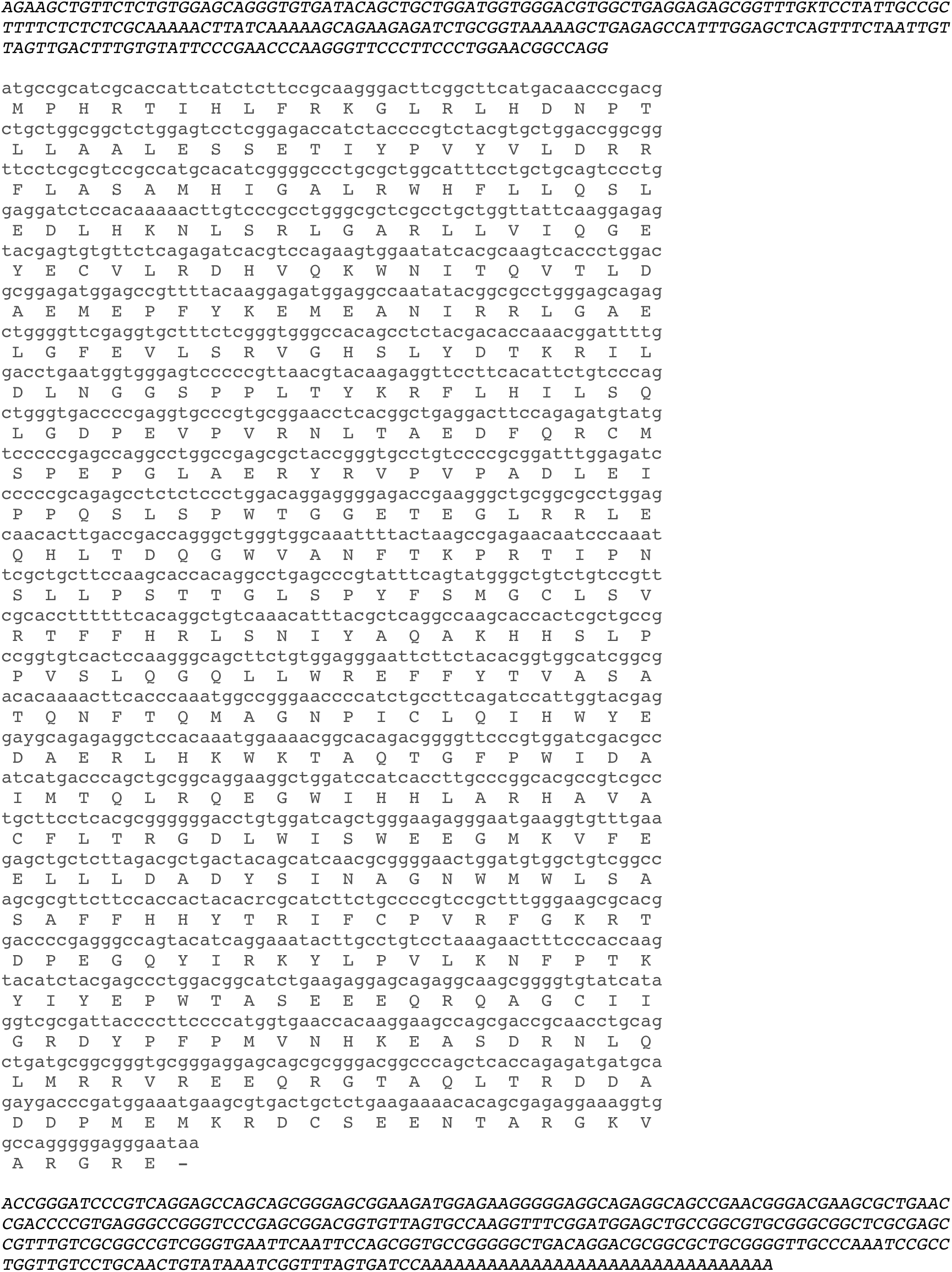
cDNA sequence and amino acid translation of *Cry4* from *Columba livia*. Rapid amplification of cDNA ends permitted cloning of the full length cDNA sequence of CRY4 from the pigeon retina. The peptide encoded is 525 amino acids in length. The 5’ and 3’ untranslated regions are shown in italiz.

**Supplementary Figure 2.**
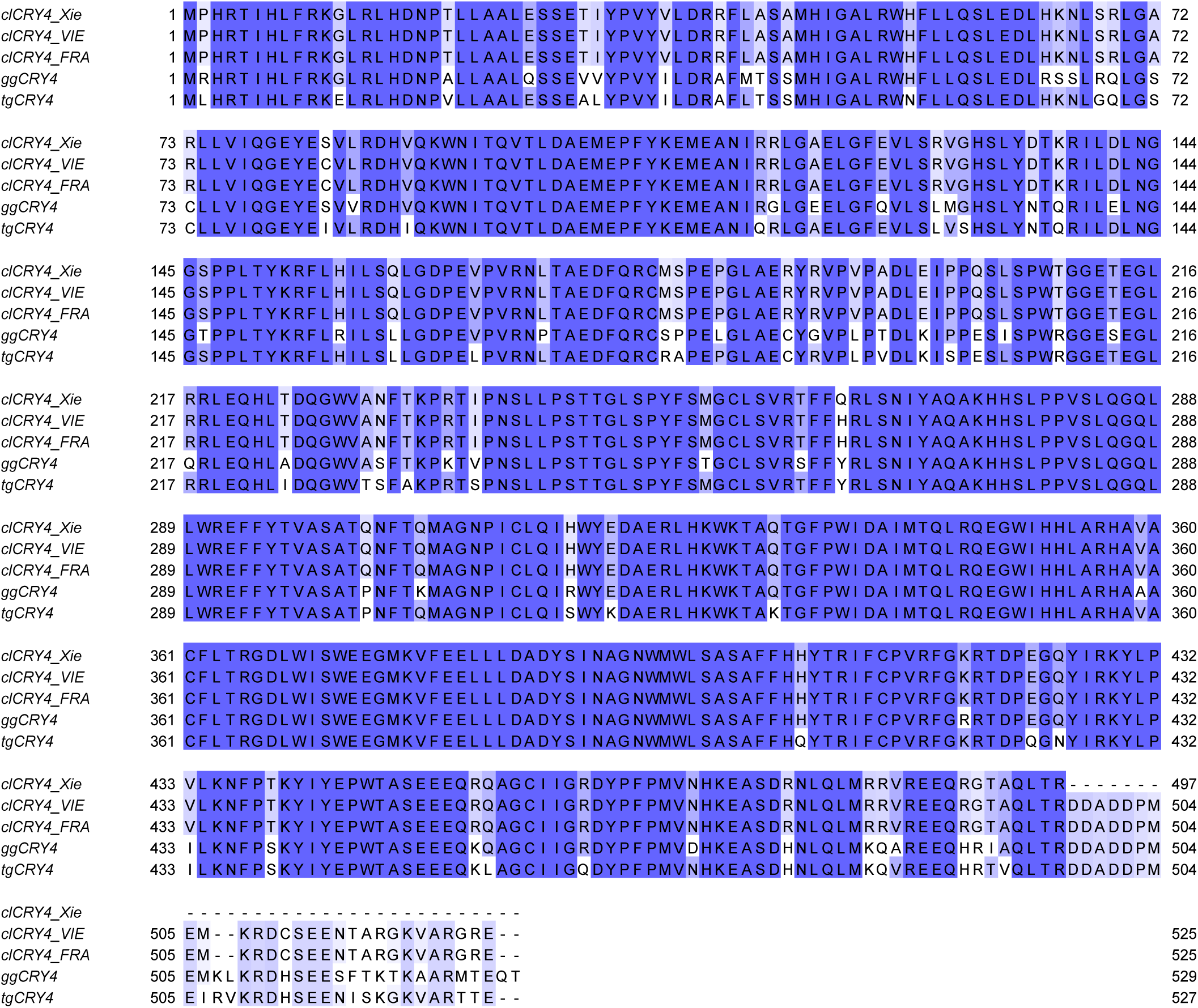
Alignment of CRY4 from different species. Protein sequences of the CRY4 peptide used by Xie and colleagues (clCRY4_XIE) which is 497 amino acids long compared to the full length CRY4 (525 amino acids) that we cloned from two different cohorts of pigeons (clCRY4_ViE, clCRY4_FRA). ggCRY4 shows the sequence of chicken CRY4 which is 529 amino acids in length, and zebra finch CRY4 which is 527 amino acids long (tgCRY4).

**Table S1.**
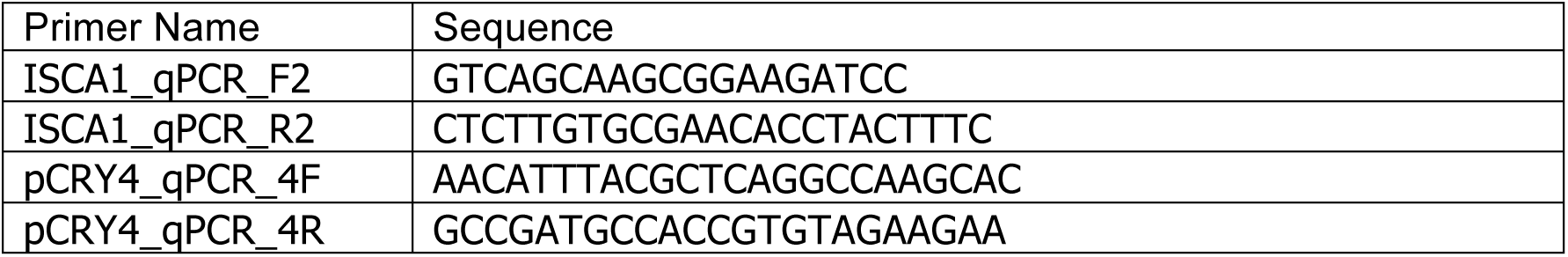
Primers used for qPCR experiments

